# Incorporating habitat selection does not account for nonrandom camera deployment in a design-based viewshed density estimator

**DOI:** 10.1101/2025.08.22.671559

**Authors:** Ehsan Moqanaki, Jennifer L. Stenglein, Glenn E. Stauffer, Daniel J. Storm, Darren Ladwig, Eli Wildey, Martha W. Zillig, Paul M. Lukacs

## Abstract

Camera trap-based abundance estimators are increasingly used for population size estimation in the absence of marked individuals. One approach is to relate animal detections to the space sampled by each camera’s viewable area, which results in viewshed density estimates that can be extrapolated to the broader sampling area to obtain abundance. The assumption is that spatial variation in local abundance corresponds to the collective viewsheds of cameras. Therefore, these design-based viewshed density estimators require that camera locations be representative of the study area. This assumption can be met with spatially balanced probability sampling, such as a generalized random tessellation stratified design. However, the random placement of cameras can be restrictive in practice. Using simulations and an empirical study, we evaluated an extension of the instantaneous sampling estimator to account for unmodeled spatial variation in local abundance using independent predictions of relative habitat use from GPS telemetry data. We applied this approach to a fenced population of white-tailed deer (*Odocoileus virginianus*), some of which were GPS collared, where timelapse photography data were collected using random and nonrandom camera placements simultaneously during two consecutive winter seasons. Our simulations showed that this approach neither reduces bias nor improves the precision of abundance estimates considerably. Specifically, we found little support that calibrating estimates based on habitat selection analysis would produce unbiased results when sampling is spatially unbalanced and focused on areas of high animal use instead of being representative of the study area. Our findings underscore the need for randomized sampling when the goal is to provide unbiased population size estimates for unmarked wildlife populations using design-based viewshed density estimators. Attempts to relax the model assumptions must be both theoretically sound and practically feasible, otherwise there would be a risk of misleading management decisions by using unreliable estimates of population size.

## 1. Introduction

Camera traps have emerged as a leading noninvasive data collection tool in ecological surveys and monitoring. A key area where camera traps have been used from their early stages of inclusion as a sampling tool is estimating population size (Buckland et al., 2023; Delisle et al., 2021). Early methods relied on capture-recapture methods where photographed individuals could be identified, and therefore individual detection histories could be constructed (Foster & Harmsen, 2012; Karanth, 1995). Although there are extensions of capture-recapture for unmarked populations (Chandler & Royle, 2013), estimating abundance in the absence of uniquely identifiable individuals is challenging. Several methods have been developed for abundance estimation of unmarked populations that we collectively refer to as camera-based abundance estimators (Moeller et al., 2023). Each method has design considerations, data requirements, and model assumptions that must be met to obtain reliable abundance estimates (Amburgey et al., 2021; Gilbert et al., 2021). Therefore, the performance of these estimators can vary substantially under different conditions.

Design-based abundance estimators such as space-to-event (STE) and instantaneous sampling (IS) attempt to relax the need to account for varying imperfect detection by using timelapse photography, accurate measures of the sampled area in front of each camera (viewshed), and random camera placement (Moeller et al., 2018). Modern camera traps can typically be programmed to take photos at fixed time intervals (timelapse) or when motion is detected. Motion-triggered photography maximizes the number of animal detections but complicates accurate measurements of viewsheds and the interpretation of detection probability (Carswell et al., 2025; Moeller et al., 2023). When cameras are programmed to take timelapse photos, the ability to detect the target wildlife when the species is present is only dependent on the ability to see them in the photos. Therefore, the detectability within the unobstructed field of view (FOV) of each camera can be considered perfect. Besides a tenable perfect detection assumption, using timelapse photography means that each sampling occasion is a snapshot moment in time and, therefore, the STE and IS estimators do not require data on animal movement (Moeller et al., 2018).

Assuming homogeneous density, animal abundance could be estimated within the camera viewsheds and then extrapolated to the entire area of interest. However, species are unlikely to be randomly distributed in space. To account for spatial heterogeneity in abundance, design-based estimators require camera locations to be representative of the sampling frame, meaning camera traps must sample the focal landscape and varying features representatively (Gilbert et al., 2021; Moeller et al., 2018, 2023). Conventional wildlife camera trapping often focuses on sampling microhabitats to maximize detections; for example, on dirt roads and game trails that many species use to move across the landscape, or landscape features such as signposts or other attractants where a target species might visit regularly (Hofmeester et al., 2019; Tourani et al., 2020). Site accessibility for practitioners may also limit representative sampling of a landscape, where sampling is biased towards sites closer to roads. There is evidence that violating the sampling representativeness assumption and a bias towards areas of high use can result in substantial overestimates of population size using design-based estimators (Amburgey et al., 2021; Tellier, 2024). Therefore, practitioners are interested in novel methods that achieve reliable estimates of population size despite nonrandom camera trapping designs.

One approach to account for sampling bias towards areas of high animal use is to incorporate auxiliary data sources that capture heterogeneity in animal space use into population density estimates. If an estimator assumes that sampling locations are random with respect to the study population, the condition can be relaxed by incorporating additional data such as GPS telemetry and accounting for the proportion of the population at every sample location in the model (Diefenbach et al., 2025; Marques et al., 2013). The idea is that by modeling the distribution of the target species relative to the sample locations, a nonrepresentative sample of the study area can be used to estimate population size reliably. An alternative strategy would be to calibrate the density estimates using auxiliary space use information to account for unmodeled spatial heterogeneity in local abundance (Henrich et al., 2025; Luo et al., 2020). If animal detections are driven primarily by the presence of a resource or habitat characteristic that could violate the representative sampling assumption, an inverse probability weighting approach can be used to adjust for sampling bias (Malchow & Hartig, 2024; Thompson, 2013). This could involve adjusting population size estimates until the model output closely matches the results from auxiliary data using an optimization method. To ensure that spatial variation within the sampled population approximates that of the target population, every sampling unit can be given a sampling weight using auxiliary information and, therefore, the probability of sampling at a given location is proportional to its weight (Malchow & Hartig, 2024). Therefore, if sampling is not representative of the study landscape, theoretically the bias and precision of an estimator can be improved by an independent estimate of relative habitat use or predicted density at camera viewsheds. However, an extensive evaluation of this approach to improve the performance of design-based viewshed density estimators is lacking. Using simulations, we evaluated an extended version of the IS estimator that incorporates predicted relative habitat use at camera locations. We apply this approach to an empirical case of white-tailed deer (*Odocoileus virginianus*) population monitoring, where timelapse photos from both random and nonrandom camera placements were used together with GPS telemetry movement data.

## 2. Materials and Methods

### 2.1 General approach

Here, we focus on the IS estimator because, unlike the STE model, viewshed-level density estimates can be obtained using the IS estimator (Moeller et al., 2018). The IS estimator uses counts of animals in the FOV of timelapse cameras that are randomly deployed across the study landscape. The model assumes geographic closure at the sampling frame level, perfect detection within the FOV of each camera, random placement of cameras, no spatial or temporal correlation in camera detections, random animal movements independent of cameras (i.e., no behavioral responses), and that there are no species misidentifications nor inaccurate counts of animals in timelapse photographs. Under these assumptions, abundance can be estimated as the total number of animal encounters at camera *j* = 1, 2, …, *J* at instantaneous sampling occasion *k* = 1, 2, …, *K* as fixed area repeat counts over space and time: 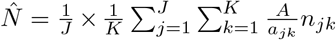. Here, *n*_*jk*_ is the count of all encounters of animals observed at a given camera and occasion across all FOV of cameras *a*_*jk*_, and *A* is the area of the sampling frame. Sampling occasions are simultaneous and instantaneous moments in time (i.e., unique date and time) when every camera has collected species group counts *n*_*jk*_ = 0, 1, 2, …, |*A*|. The sampling variance of 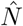 is estimated via the delta method, and confidence intervals (CIs) are obtained via bootstrapping (Moeller et al., 2018).

Boyce and McDonald (1999) described how to estimate probability of use from resource selection functions (RSFs) and availability of habitat covariates. For a vector of spatial covariates **x**_**i**_ at locations *i* = 1, 2, …, *I*, probability of use is 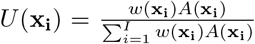, where *w*(.) and *A*(.) are the RSF and area of habitat types, respectively. Here, an RSF could be any function that is proportional to probability of use of a resource unit (Fieberg et al., 2021). If availability of habitat types and population size remain constant under the condition in which the RSF was derived, then **x**_**i**_ = *a*_*jk*_. The relationship between camera-specific density estimates obtained by the IS estimator 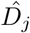 and predicted habitat use at cameras’ viewshed areas *U* (**x**_**i**_), rescaled between 0 and 1, can be quantified using a linear model: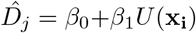, where *β*_0_ and *β*_1_ are the intercept and slope, respectively. Then, a calibrated abundance estimate is as follows: 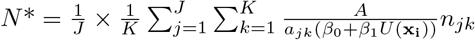.

Since viewshed-level density estimates by the IS estimator can be right-skewed and have excess zeros – i.e., nonnegative data with clumping at zero, we evaluated two additional approaches to obtain a calibrated estimate of abundance by: (1) fitting a Tweedie generalized linear model (GLM) using different combinations of the family and link function (Dunn, 2022); and (2) using 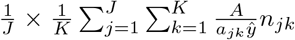, where 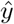 is the mean of predicted relative habitat use at camera locations. We used this framework to prepare and analyze simulated and empirical data in R 4.4.2 (R Core Team, 2024), and we used `spaceNtime` (Moeller & Lukacs, 2022) to construct detection histories and fit the IS estimator (Supporting Information).

### 2.2 Simulations

We used simulations to evaluate conventional and calibrated IS estimates in situations of homogeneous and spatially heterogeneous distribution of individuals where sampling design was either spatially balanced and representative of the study landscape or spatially biased towards areas of high use (Fig. 1). We used our study of white-tailed deer in Wisconsin to guide our simulation scenarios and settings (Supporting Information). White-tailed deer are semi-gregarious animals that form both family groups and mixed foraging groups with substantial variations in their group sizes and movement behavior over time and across different landscapes (McShea, 2012). We evaluated four simulation scenarios illustrated in Fig. 1, where the population consisted of (1) both solitary individuals and family groups of two to six individuals with minimal individual heterogeneity in their movement patterns; or (2) solitary movements with substantial individual variation in space use (Supporting Information).

**Figure 1.**
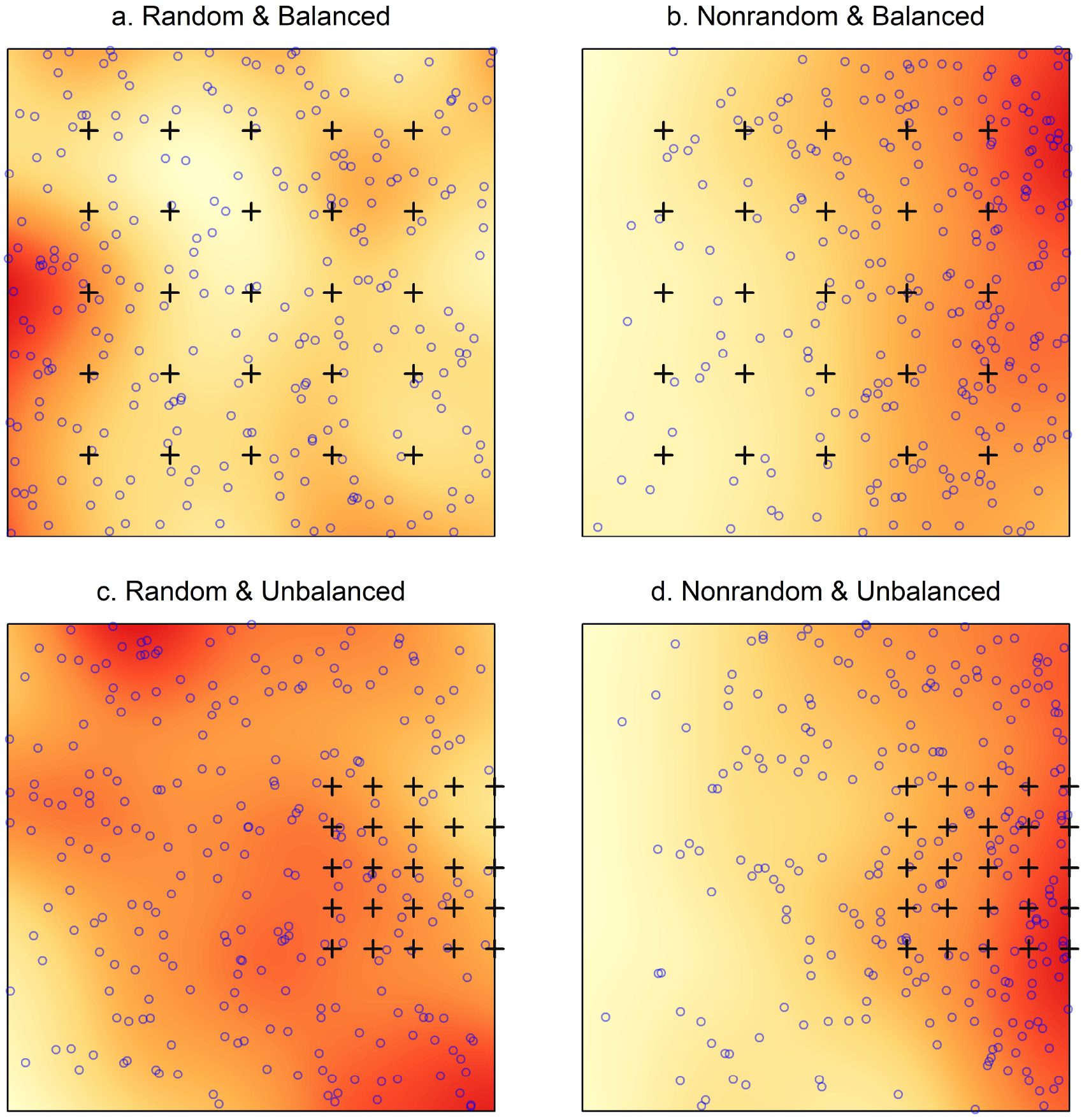
Illustration of four simulated scenarios of random and nonrandom distribution of individuals (blue circles), where the camera trapping design (black crosses) was either spatially balanced and representative of the 6000 × 6000-distance unit (du) study landscape or was spatially unbalanced and biased towards a subset of the study area. The underlying surface represents density of starting points of simulated individual movements at 20-du pixel resolution as a measure of relative habitat use, where warmer colors show higher density of starting points.

For all simulations, we used a 6 000 × 6 000-distance unit (du) square grid as the study area (*A* = 36 × 10^6^), and we simulated 252 starting points of individual movements as the total population size *N* . In scenarios of random distribution of the simulated population, starting point locations were randomly distributed across the study area without introducing landscape heterogeneity (Fig. 1a,c). For scenarios of nonrandom, spatially heterogeneous, distribution of individuals, we considered population density decreases linearly from east to west by applying a function to randomly place 10 times more starting point locations at the east end of the study area compared to the west end (Fig. 1b,d). We chose this scenario to represent situations where, because of a suitable habitat type, resource, or feature of high use such as trails, individuals are clumped in a region at a given sampling time, representing strong habitat preferences. The spatially balanced sampling design specified a regular grid of 5 × 5 camera traps with 1 000 du inter-camera spacing (Fig. 1a,b). The spatially biased design specifically targeted the east side of the study landscape (500 du camera spacing; Fig. 1c,d), thus violating the representative sampling assumption in cases where animal distributions were biased towards cameras (Fig. 1d). We defined the FOV of each camera using a circular sector of 20-du radius as the maximum viewable distance from the camera and 35^°^ viewable angle. We assumed perfect detection within camera FOVs.

From each starting point, we simulated movements as discrete-time, biased, correlated random walks (Supporting Information). We defined duration of timesteps as one minute and ran the movement simulations for 30 days of 12-hour activity. We allowed substantial home range overlaps during data generation, but associations in movement were limited to scenarios with group sizes larger than one. We restricted animal movements within the study extent to keep population density constant at any point in time. Thus, all scenarios represented geographically and demographically closed populations with densities of 7 × 10^−6^ individuals/du^2^. We sampled animals’ trajectories overlapping with each camera’s viewable area at 15-minute time intervals (0, 15, 30, and 45 minutes of each hour) to be consistent with data collection using timelapse photography. A timelapse detection was collected only if one or more simulated animals were in the FOV of a camera in each timelapse instant.

We considered five alternative predictions of relative habitat use where the source of spatial variation in abundance was either known and correctly estimated or was misspecified (Fig. S1). Continuous relative habitat use surfaces were scaled to have values between 0 and 1. Discrete relative habitat use surfaces were created using continuous layers by selecting median values as cut offs and assigning values of 1 or 0.1 to habitat cells. In total, for each simulation scenario we compared 16 estimates of abundance by the standard IS estimator (*n* = 1) and after calibration (*n* = 15) using five alternative realizations of relative habitat use and three calibration approaches of fitting a linear model, a Tweedie model, or using average predicted habitat use values. We repeated the data simulation 200 per scenario, resulting in a total of 800 simulated timelapse photography data sets per movement scenario, or 25 600 model runs including 1 600 IS and 24 000 calibrated abundance estimates. We report mean and 95% CI of abundance estimates. We evaluated model performances in each group of scenarios based on mean estimates of abundance in terms of (1) relative bias (RB) from true population size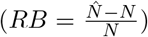, (2) relative stan-dard error 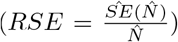 and (3) the confidence interval width (CIW; difference between the 97.5^th^ and 2.5th percentiles of the mean) as two measures of precision, and (4) the nominal coverage of the 95% CI of true population size. Unless specified otherwise, we report all results based on simulations with mixed-group movements and refer the reader to Supporting Information for additional scenarios.

### 2.3 Case study

We applied the calibration approach to monthly abundance estimates of a free-ranging, fenced, population of white-tailed deer during January-March 2022 and 2023 in Sandhill Wildlife Area (SWA), Wisconsin, USA (Fig. S2). Helicopter counts suggested a minimum population size of between 143 and 153 deer in March 2022, and between 207 and 236 deer during February-March 2023. For habitat analysis using GPS telemetry data, we sampled a total of 27 individual deer of different sex and age classes (Table S1). Detailed descriptions of data collection, preparation, and analysis are provided in Supporting Information. We constructed winter-specific RSFs to quantify the link between locations of individual deer and important environmental features (Fieberg et al., 2021). We identified winter as December 1 to April 30 based on knowledge of deer ecology in SWA. We considered five habitat covariates at 20 × 20 m resolutions that might have influenced deer habitat-selection patterns (Fig. S3 and Table S2). We fitted a generalized linear mixed model with additive effects of each covariate and the random effect of individual-year using `lme4` (Bates et al., 2015). Relative habitat use was calculated from predicted RSF values as described above.

We installed 21 randomly-placed and 21 trail-focused (nonrandom) cameras across the study area to collect timelapse data for abundance estimation (Fig. S2). Cameras used infrared low-glow LED flash technology (Bushnell Outdoor Products, Kansas, USA). We placed cameras approximately 80 cm above the ground, oriented north, and programmed to take one timelapse photograph per 15 minutes. We used timelapse images from two consecutive years in January-March 2022 and 2023 to coincide with aerial surveys that provide rough deer population size estimates. We calculated camera-specific FOVs using maximum viewable angle and distance derived from our own experiments and manufacturers’ specifications (Supporting Information), where detection was assumed to be perfect. We considered 84% of SWA (*A* = 31.2 km^2^) as deer habitat for abundance estimates (Figs. S2 and S4). We compared estimates by the IS estimator with three calibrated estimates against the total counts from aerial surveys. For comparison, we also reported estimates by the STE model, which has similar data requirements and assumptions (Moeller et al., 2018), totaling five estimates per sampling design (random vs. nonrandom). Data and R code for model fitting will be provided in Supporting Information.

## 3. Results

### 3.1 Simulations

In most instances of spatially heterogeneous abundance and spatially unbalanced sampling (Fig. 1d), we observed a spike in individuals detected, total timelapse detections, maximum number of individuals in timelapse detections, and cameras with at least one detection compared to the other three scenarios (Fig. S5). The IS estimator was robust to spatial variation in local abundance, provided the camera trapping design was representative of the study landscape and that the full range of population density were sampled in a spatially balanced manner (Fig. 2a,b and Fig. S6a,b). The IS estimator exhibited noticeable negative bias 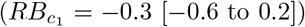 in the scenario of homogeneous population density with a spatially unbalanced sampling design (Fig. 2c), and pronounced positive bias 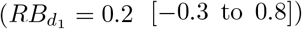 in the scenario with spatial heterogeneity and cameras placed in high use areas (Fig. 2d). We observed comparable RSE and CIW of the IS estimator across the simulation scenarios, often with good precision (*RSE* ≈ 0.2). Nonetheless, coverage probability was below nominal in scenarios of spatially unbalanced sampling (Fig. 2c,d), declining from 0.94 to as low as 0.73.

**Figure 2.**
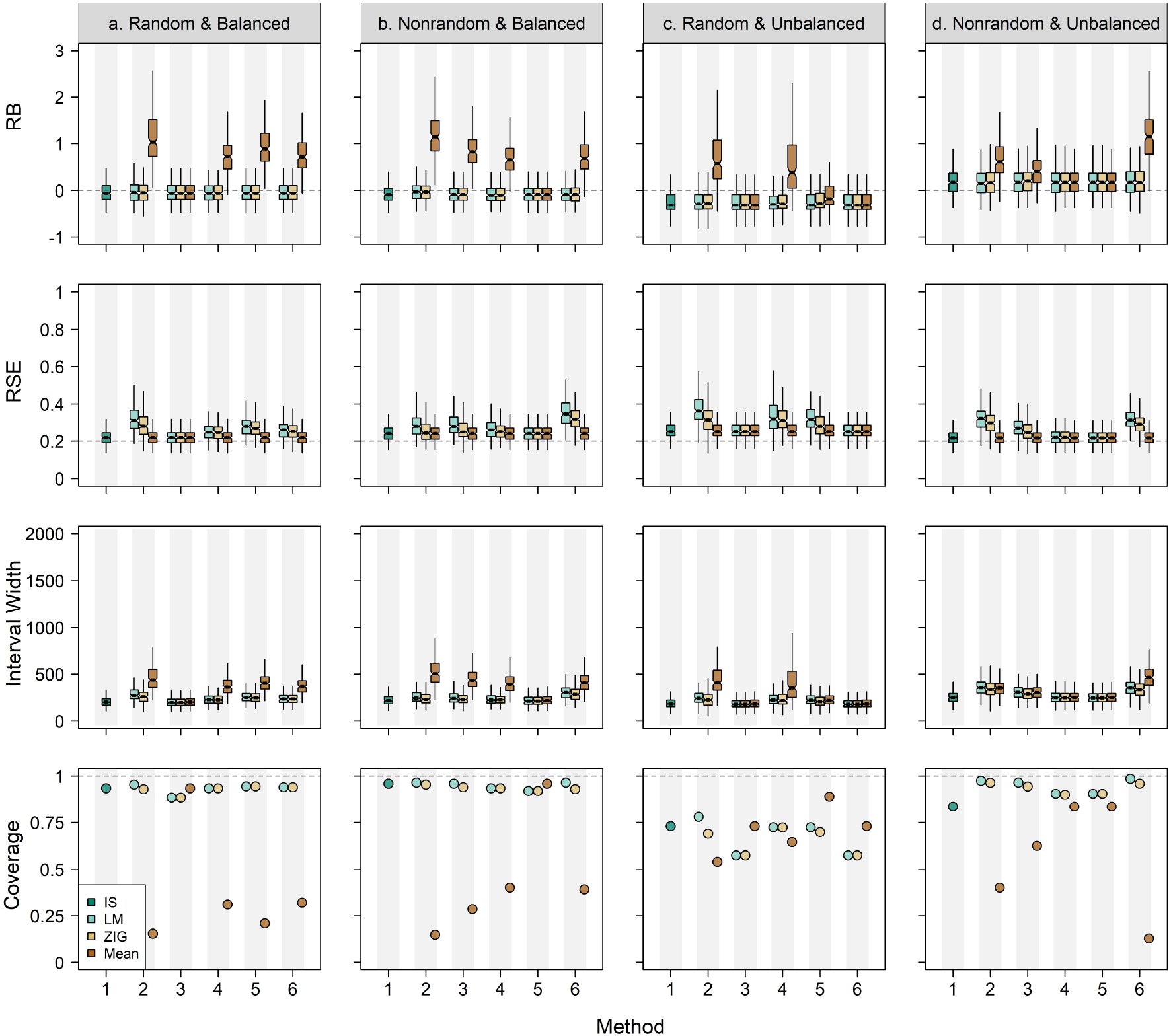
Relative bias (RB), relative standard error (RSE), confidence interval width, and coverage probability of the 95% confidence intervals of true population size estimated by the instantaneous sampling (IS) estimator (Method 1) compared to calibration methods using five alternative predictions of relative habitat use (Method 2-6; Fig. S1). We tested three calibration approaches using a linear predictor (LM), a Tweedie model (ZIG), and mean of predicted relative use values at camera locations (Mean).

Using calibration methods, we observed minimal gains in incorporating space use information into the abundance estimates regardless of whether there was spatial variation in population density and camera trapping designs. Patterns in RB were qualitatively comparable within scenarios regardless of incorporating habitat use information and fitting either a linear predictor or a Tweedie GLM. However, we detected substantial positive bias up to 110% by calibrating IS estimates using average relative use values at camera locations, even if an accurate prediction was incorporated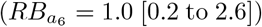. The only noticeable exception was in scenario of homogeneous population density with spatially unbalanced sampling design (Fig. 2c), where calibrated density estimates using mean values of an inaccurate index of relative habitat use were on average less biased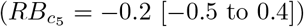.

Compared to the conventional IS estimator, RSE was almost always lower only when mean of relative use values were used for calibrating the estimates, although the pattern appeared inconsistent, and the aforementioned pattern was not limited to situations where an accurate realization of relative habitat use was included. We detected considerable loss of precision in many situations, even if density estimates were calibrated with accurate estimates of relative use (Method 2). The only apparent gain was in coverage probability, which increased to up to 1 by calibration using either a linear predictor or Tweedie GLM model in the scenario of spatially heterogeneous density with spatially unbalanced sampling (Fig. 2d). However, these results often represented an excessively wide interval, rather than actual improvements in estimates of abundance. Calibration by mean values of relative use reduced coverage probability substantially in most situations, to as low as 0.13.

### 3.2 Case study

Using timelapse data from randomly placed cameras, white-tailed deer abundance estimates by conventional STE and IS estimators almost always included the range of total counts by aerial surveys (Fig. 3). However, by using data from trail-focused cameras IS estimates were on average between 1.8 and 3.9 (2022) and 0.7 and 13.0 (2023) times higher than IS estimates using data from randomly-placed cameras, a pattern consistent with STE estimates. We noticed a drop in the 2023 abundance estimates, presumably because of camera failures. After incorporating predicted relative habitat use (Figs. S7-S11, Tables S3-S4), we observed inconsistent patterns in the resulting estimates of abundance. Although there were a few instances where calibrated IS estimates were closer to aerial counts than were conventional IS estimates, mean of abundance estimates remained relatively unchanged or deviated more from reference total counts in most instances (Fig. 3). Calibration almost always increased CIW considerably.

**Figure 3.**
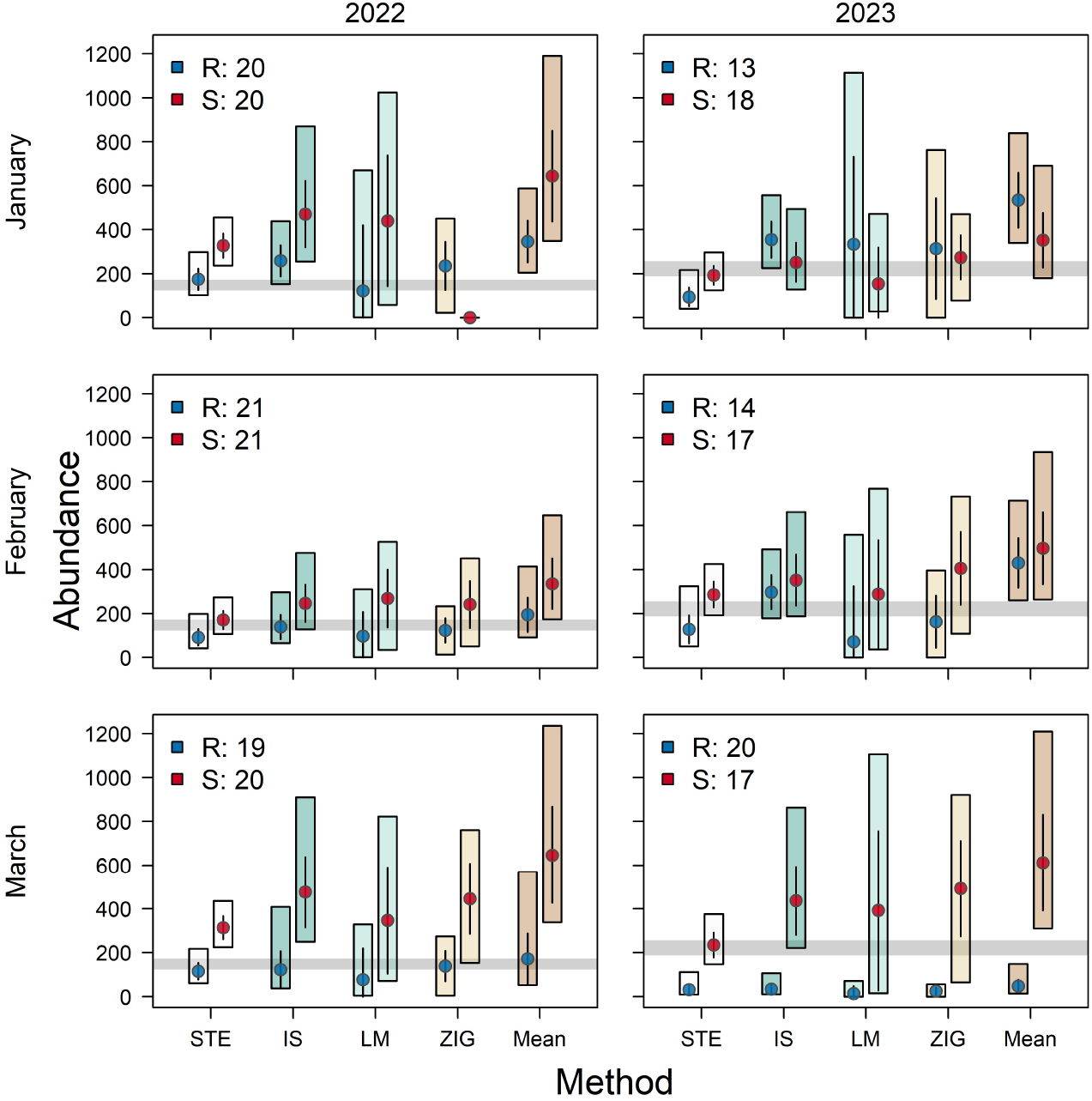
White-tailed deer (*Odocoileus virginianus*) abundance estimates for Sandhill Wildlife Area, Wisconsin, by all methods using timelapse camera-trap data from a total of 21 random (R) and 21 trail-focused cameras (S) during January-March 2022 and 2023: space to event (STE) model, instantaneous sampling (IS) estimator, and calibrated density estimates using predicted relative habitat use at camera locations via a linear predictor (LM), a Tweedie model (ZIG), or mean of predicted values (Mean). Colored bars show the 95% confidence interval, and the circles and lines represent mean and standard error of the estimates, respectively. Dark gray lines represent total counts based on multiple aerial surveys via helicopter per winter season as a rough estimate for population size during the sampling period.

## 4. Discussion

We demonstrated a new approach to incorporate auxiliary space-use information into the design-based IS estimator of abundance. We identified situations where this approach could improve estimates of abundance. However, we found little support for relaxing the representative sampling assumption of the IS estimator.

### 4.1 Incorporating space-use information into camera-based population size estimation

Linking spatial heterogeneity in local abundance to space use is common in the camera trapping literature, because animal detectability can be a direct result of varying space use (Hofmeester et al., 2019; Tourani et al., 2020). Different methods have been proposed to incorporate additional space use information into camera-based population size estimation (Luo et al., 2020). Henrich et al. (2025) described a rather similar approach to calibrate estimates by the random encounter model (Rowcliffe et al., 2008) and camera-trap distance sampling (Howe et al., 2017) using realized relative population density obtained from an independent spatial capture-recapture (SCR) analysis of noninvasive genetic sampling and habitat analysis of GPS telemetry data. Their work can be compared to one of our models, where density estimates were calibrated using the mean values of relative use at camera locations. This inverse calibration assumes that average population size can be reliably estimated from a small sample of the study population with the condition that the relative value of sampling units to true population size is known and can be used in a weighted average (Henrich et al., 2025; Malchow & Hartig, 2024). Here, uncertainty in weights drives uncertainty in estimated population size; assuming that true weights are fully known via auxiliary space use information in an ideal scenario, average population size can be estimated perfectly. Although Henrich et al. (2025) suggested this approach could be useful for calibrating two model-based estimators, particularly with incorporating independent SCR results, we found the least reliable results using this method. The contrasting finding can be explained not only by differences in model fitting, simulated scenarios, and study system, but also in fundamental difference between inference from model-based and design-based approaches. The former relies on distributional assumptions about the underlying data-generating stochastic process, whereas the latter, including the IS estimator, relies on random sampling – i.e., randomly assigning some population units to be in the sample (Dumelle et al., 2022).

### 4.2 In which situations could the calibration approach be useful?

We detected some gains by calibrating the IS estimates with accurate relative use values where data were generated from spatially balanced sampling. Likewise, we observed indications for such possible improvements during our case study. Our simulations were influenced by our study of a fenced white-tailed deer population, a habitat generalist ungulate species restricted to a predominantly suitable landscape where we did not detect substantial spatial variation in habitat use across the area considered to be deer habitat. In situations where there is a strong habitat preference over patchy resources, for example, in monitoring a habitat specialist in a heterogeneous landscape or a habitat with scarce water or food in which a subset of cameras is installed to monitor these resources, we expect that our calibration approach could be more useful. However, we note that these targeted localized resources can trigger behavioral responses of wildlife to camera locations, and that the calibration approach might not be sufficient to relax the design requirement of camera locations being random relative to animal movement, which is a requirement for several camera-based estimators but not the unmarked SCR and site structured (*N* -mixture and Royle-Nichols) models (Gilbert et al., 2021).

Further simulation studies and empirical comparisons could be helpful in evaluating tradeoffs in using calibration methods with other camera-based estimators. In cases where motion-triggered data are collected, this type of data can be used for abundance estimation using the design-based STE model (Elbroch et al., 2024; Lyet et al., 2024). Similar approaches can be followed by using the IS estimator with motion-triggered data to evaluate a wider range of situations encountered in the field for monitoring unmarked wildlife populations. These calibration methods can be also extended to any other habitat selection analysis that incorporates movement characteristics to produce spatially explicit habitat use such as step selection functions (Fieberg et al., 2021), as well as density surface modeling (Miller et al., 2013; Tourani, 2022).

### 4.3 Practical and methodological limitations

Our results confirm that the conventional IS estimator performs relatively poorly where population density is highly variable across the study area, sampling is not representative, and animal movements are biased towards sampling locations. Although we observed some improvements in abundance estimates, our calibration methods did not decrease bias or improve precision when sampling design was spatially unbalanced and nonrepresentative. We expect that in cases where reasonably high camera density is not used for monitoring (Luo et al., 2020) and there is little variation in animal use intensity within the FOV of cameras, these calibration methods would perform poorly.

The utility of this approach is also limited to situations where high-quality auxiliary information exists, and reliable inferences are made about underlying habitat selection or population density to calibrate abundance estimates. Collection of additional data sources by GPS telemetry or noninvasive DNA sampling can be labor intensive and costly, jeopardizing the main advantages of camera-based abundance estimators of unmarked wildlife (Gilbert et al., 2021; Wearn & Glover-Kapfer, 2019). In addition, habitat selection analyzes that are suitable for the calibration approach would often require capturing and fitting a subset of the study population with GPS collars, collection of fine-resolution movement data, and availability of viewshed-level determinants to quantify habitat selection, which could make the method inapplicable to large areas. In our study, we hypothesized that linear movement paths influenced deer habitat selection and, in turn, their detections within the FOV of cameras substantially, supported by higher numbers of timelapse detections using on-trail vs. off-trail cameras (Fig. S2). However, we were unable to map and include all such spatially fine features across the study area in our habitat selection analysis. Quantifying how species perceive and respond to fine-scale habitat characteristics using relatively crude landcover covariates is unlikely to be very informative for calibration, especially for habitat generalists.

Independent, spatially explicit estimates of abundance such as those suggested by Henrich et al. (2025) can be particularly useful, but they are also restrictive, represent only a snapshot of population density, and are more suitable for smaller areas, although several large-scale applications exist (Tourani, 2022). We noticed substantial seasonal changes in white-tailed deer space use in our study, especially during the mating period in late fall. Predictions of habitat use or population density must account for these temporal variations to be useful for calibration of population size estimates. Practitioners need to select reasonably short sampling durations with relatively stable areas of use intensity that match the camera trapping period, particularly a period with no major habitat disturbance or changes in animal behavior (e.g., breeding and hunting seasons). Predictions at a different time can only be used if selection intensity or population density has not changed. However, this assumption would be difficult to support without tracking changes over time in the first place. In addition, incorporating uncertainty in the estimates of relative use or population density into the calibration methods is challenging. In a Bayesian framework, methods developed for integrated modeling (Schaub & Abadi, 2011) can handle the need to propagate the error from RSFs or spatially explicit population size models to calibrated abundance estimates.

### 4.4 Implications for camera-based monitoring of unmarked wildlife

Inverse calibration has potential to improve camera-based abundance estimates of unmarked populations when spatial heterogeneity in abundance remains partially accounted for. However, we demonstrated that such gains might come with considerable cost and could be limited in practice. Poor applications of this approach can introduce substantial bias and result in loss of precision of the estimators. Using the suite of design-based density estimators evaluated, we believe that meeting the representative sampling assumption by adopting an efficient randomized sampling approach, such as stratification or adaptive sampling, should be the priority for study design and data collection. We caution against simplified approaches to scale population size estimates based on sampling a small subset of heterogeneous landscapes, unless an appropriate model-based solution is incorporated to account for different sources of variable detectability resulting from spatially unbalanced sampling.

## Acknowledgments

We thank the staff and volunteers from the Wisconsin Department of Natural Resources for their support with field logistics and photo classification during different stages of this project, including D. MacFarland, J. Winiarski, S. Lee, G. Freeman, A. Schneider, C. Milestone, Sandhill Wildlife Area staff, and Snapshot Wisconsin volunteers. E. Coen-Pesch, E. Hachfeld, and M. Cook assisted in classifying camera trap photos. Thanks to M. Tourani for helpful suggestions and making us aware of the unpublished study by Henrich et al. Any use of trade, firm, or product names is for descriptive purposes only and does not imply endorsement by the U.S. Government.

## Author contributions

EM, PL, JS, and GS conceived the idea and designed methodology; JS, DJS, DL, GS, and EW collected and supplied the empirical data; EM, JS, MZ, and EW prepared the data; EM analyzed the data and led the writing of the manuscript. All authors contributed critically to the drafts and gave final approval for publication.

## Ethical Statement

All aspects of fieldwork, including captures and handling of deer, were in accordance with protocols approved by the Wisconsin Department of Natural Resources within guidelines of the American Society of Mammalogists (Sikes and The Animal Care and Use Committee of the American Society of Mammalogists, 2019).

## Data Availability Statement

The empirical data set used in this study is available from the Wisconsin Department of Natural Resources, but restrictions may apply to the availability of these data. Request of these data can be made to. Data and R code exemplifying the analysis will be deposited in Github upon acceptance.

## Supporting Information

Additional supporting information may be found online in the Supporting Information section at the end of the article.

